# IgY antibodies against Ebola virus possess post-exposure protection and excellent thermostability

**DOI:** 10.1101/2020.05.21.108159

**Authors:** Yuan Zhang, Yanqiu Wei, Yunlong Li, Xuan Wang, Yang Liu, Deyu Tian, Xiaojuan Jia, Rui Gong, Wenjun Liu, Limin Yang

**Affiliations:** Key Laboratory of Pathogenic Microbiology and Immunology, Institute of Microbiology, Chinese Academy of Sciences, Beijing, China; Anhui University, Hefei, Anhui, China; Institute of Microbiology, Center for Biosafety Mega-Science, Chinese Academy of Sciences, Beijing, China; CAS Key Laboratory of Special Pathogens and Biosafety, Wuhan Institute of Virology, Center for Biosafety Mega-Science, Chinese Academy of Sciences, Wuhan, Hubei, China; The Biological Safety level-3 (BSL-3) Laboratory of Institute of Microbiology, Chinese Academy of Sciences, Beijing, China; Shenyang Agricultural University, Shenyang, China

## Abstract

Ebola virus (EBOV) is the most virulent pathogens that cause hemorrhagic fever with high mortality rates in humans and nonhuman primates. The postexposure antibody therapies to prevent EBOV infection are considered efficient. However, due to the poor thermal stability of mammalian antibody, their application in the tropics has been limited. Here, we developed a thermostable therapeutic antibody against EBOV based on chicken immunoglobulin Y (IgY). The IgY antibodies demonstrated excellent thermal stability, which retained their neutralizing activity at 25°C for one year, in contrast to conventional polyclonal or monoclonal antibodies (MAbs). We immunized laying hens with a variety of EBOV vaccine candidates and confirmed that VSV Δ G/EBOVGP encoding the EBOV glycoprotein could induce high titer neutralizing antibodies against EBOV. The therapeutic efficacy of immune IgY antibodies *in vivo* was evaluated in the newborn Balb/c mice model. Lethal dose of virus challenged mice were treated 2 or 24 h post-infection with different doses of anti-EBOV IgY. The group receiving a high dose of 10^6^ NAU/kg (neutralizing antibody units/kilogram) achieved complete protection with no signs of disease, while the low-dose group was only partially protected. In contrast, all mice receiving naïve IgY died within 10 days. In conclusion, the anti-EBOV IgY exhibits excellent thermostability and protective efficacy, and it is very promising to be developed as alternative therapeutic entities.

**Author Summary:** Although several Ebola virus therapeutic antibodies have been reported in recent years, however, due to the poor thermal stability of mammalian antibody, their application in tropical endemic areas has been limited. We developed a highly thermostable therapeutic antibody against EBOV based on chicken immunoglobulin Y (IgY). The IgY antibodies demonstrated excellent thermal stability, which retained their neutralizing activity at 25°C for one year. The newborn mice receiving passive transfer of IgY achieved complete protection against a lethal dose of virus challenge indicating that the anti-EBOV IgY provides a promising countermeasure to solve the current clinical application problems of Ebola antibody-based treatments in Africa.

## Introduction

Ebola virus (EBOV) belongs to the Filoviridae family and the known cause of severe hemorrhagic fever in humans and nonhuman primates (NHPs). Since the epidemic of Zaire (now the Democratic Republic of the Congo, DRC) and Sudan in 1976, intermittent local epidemics persist in Africa, with a case-fatality rate of 25-90%. The most severe Ebola outbreak to date occurred in West Africa from 2014 to 2016, more than 28 000 people infected, and 11 323 patients died. Occasionally, cases have been exported into other countries through travel. The ongoing outbreak in the DRC is the second-largest Ebola epidemic on record, with 2 279 lives lost and 3 462 confirmed infections since August 2018, which prompted WHO to declare this epidemic a public health emergency of international concern. Pandemic potential, high mortality, high infectivity, and lack of preventive and therapeutic approaches make EBOV a Class A pathogen that seriously threatens public health.

The intermittent and continuous outbreak of Ebola disease (EVD) poses a challenge for public health, which promoted research on vaccines and antiviral drugs. The most effective strategy to prevent and treat EVD is antibody immunotherapy [1–5]. However, antibody treatment was not optimistic at the beginning, although antibody KZ52 had high neutralizing activity but failed to play a role in NHPs [6]. Until 2012, it was discovered that convalescent plasma can protect NHPs from lethal dose of EBOV infection, and antibody therapy strategy rekindled researchers’ interest [7]. The researchers tried to mix multiple MAbs against EBOV glycoprotein (GP) to form ZMapp (2G4, 4G7, 13C6), a new antibody combination based on cocktail therapy. It was used to treat seven infected persons, and 5 of them survived. ZMapp has an excellent protective effect in NHPs and patients and has greatly extended the window period of administration. The emergence of ZMapp has brought the peak of antibody treatment research [8]. Nowadays, antibody therapy becomes a promising approach to control EVD. Recently, anti-EBOV equine sera and ovine sera were confirmed that can protects rodent models against EBOV challenge [9–11]. These studies suggest that EBOV-specific antibodies from different species are expected to be developed as therapeutic agents.

Although EBOV therapeutic antibody has inspiring application prospects, it still has some application limitations. Firstly, the limited availability and security of convalescent plasma have hampered its global application. Secondly, MAbs have long preparation cycles and high production costs. Also, they all need strict transportation and storage conditions. However, the Ebola outbreak mainly occurred in the hot and less wealthy African regions, thus limiting the current application of antibodies. Poultry-derived IgY provides an alternative strategy for producing safe and inexpensive antibodies. IgY antibodies, the predominant serum immunoglobulin in birds, reptiles, and amphibians, are transferred from the serum of females to the egg yolk [12], where they offer passive immunity to embryos and neonates. As a potential therapeutic antibody, IgY has lots of advantages over mammalian IgG due to its structural and immunological properties [13]. It possesses excellent stability under various physicochemical conditions and lower manufacturing costs. It cannot bind to mammalian Fc receptors or complement components [14], which can avoid potential antibody-dependent infection enhancement (ADE) and adverse immune reactions. Eggs can be used to produce large amounts of yolk antibodies quickly and do not cause harm to animals. In recent years, the antibody therapy strategy based on IgY has been widely recognized, and many promising results have been reported in influenza virus [15–17], dengue virus [18], zika virus [19], hantavirus [20], severe acute respiratory syndrome [21], rotavirus and norovirus [22], etc. Previous studies have reported multiple Ebola vaccine candidates based on different platforms. These candidates exhibit different immunogenicity. To prepare high-immunity IgY antibodies, we need to screen for immunogens that can induce laying hens to produce the most robust humoral immune response.

This study aimed to produce an anti-EBOV IgY antibody and assess its antiviral efficiency. The IgY antibody was obtained from laying hens immunized with recombinant vesicular stomatitis virus vector encoding EBOV GP (VSV Δ G/EBOVGP). The protecting efficiency of the resulting IgY antibody against EBOV was evaluated by Enzyme-linked immunosorbent assay (ELISA), pseudotyped virus neutralization assay, and a mouse challenge model. The results show that the IgY antibody has excellent thermal stability and achieve protection against lethal challenge in newborn mice. Our results suggest that the potent IgY warrant further development as prophylactic and therapeutic reagents for EVD.

## Results

### Preparation of immunogens

Vaccine-elicited neutralizing antibodies (NAbs) are associated with protection against Filoviridae family mediated disease. In order to obtain the most potent anti-EBOV antibody, we prepared several EBOV immunogens based on multiple different platforms, including DNA vaccine (pCAGGS/EBOVGP), recombinant protein (rEBOVGP) or virus-like particle (EBOV-VLP) subunit vaccines, and two viral vector vaccines (VSV Δ G/EBOVGP, Ad5/EBOVGP). Western blot confirmed that these immunogens could express or contain EBOV GP that can induce NAbs in animals (Fig 1). Due to the differences in humoral immune responses induced by different vaccines, we need to screen for the most suitable immunogen for IgY antibody production.

**Fig 1.**
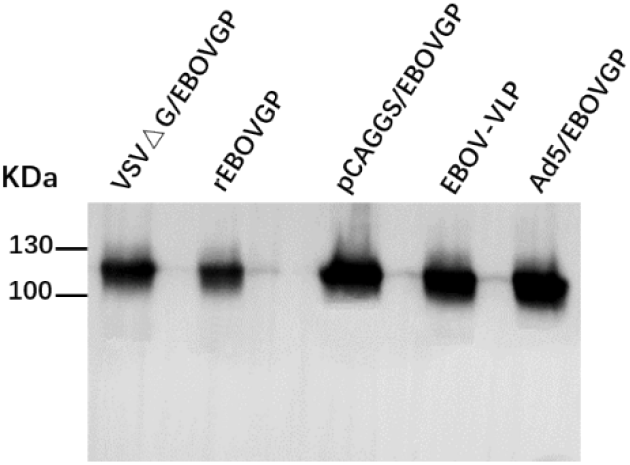
Western blot analysis of EBOV immunogens. Recombinant protein, vector immunogens and 293T cell lysates transfected with plasmid expressing EBOV GP were separated by SDS– PAGE under reducing conditions, transferred to PVDF membrane, and detected using EBOV GP1-specific MAb.

### Immunogens elicited high titer of anti-EBOV IgY

To obtain high titer EBOV NAbs, 5-month-old laying hens were vaccinated with five different immunogens, including 10^3^ or 10^4^ TCID_50_ VSVΔG/EBOVGP, 100 μg rEBOVGP, 100 μg pCAGGS/EBOVGP, 10 μg EBOV-VLP, and 10^11^ virus particles (vp) Ad5/EBOVGP (Fig 2a). Thirty-five laying hens were randomly divided into seven groups, which were immunized four times with each immunogen or PBS control intramuscularly (i.m.) at a 14-day interval. Eggs were collected at 0, 2, 4, 6, 8 weeks, and IgY antibodies were purified from egg yolk for ELISA and NAbs test. Both titers in all groups were gradually increased after the first immunization. The results showed that all immunogens except DNA vaccine induced potent Gp-specific ELISA antibodies (Fig 2b). For the NAbs, the geometric mean titers (GMTs) in 10^4^ TCID_50_ VSVΔ G/EBOVGP group reached 1:3650 (VSV pseudoneutralisation, VSV-PsN) and 1:320 (lentiviral vectors pseudoneutralisation, LVV-PsN) after the third boost, which significantly higher than other groups (Fig 2c-d). VSVΔG/EBOVGP induced the strongest NAb titers than the other immunogens. In contrast, the GP-specific ELISA and NAb titers were not detected in the negative control group. Besides, to determine the correlation between the immune dosage and antibody level, two dosages of VSVΔG/EBOVGP were used to immunize laying hens. 10^4^ TCID_50_ VSVΔ G/EBOVGP induced stronger NAbs titer compared to 10^3^ TCID_50_ VSVΔ G/EBOVGP, suggesting that the immunization dose is crucial for obtaining higher titers of NAbs. In order to exclude the possible interference of the antibody generated by the VSV vector backbone, we used two PsN assays, VSV-PsN and LVV-PsN, and the results showed that the two assays obtained consistent results.

**Fig 2.**
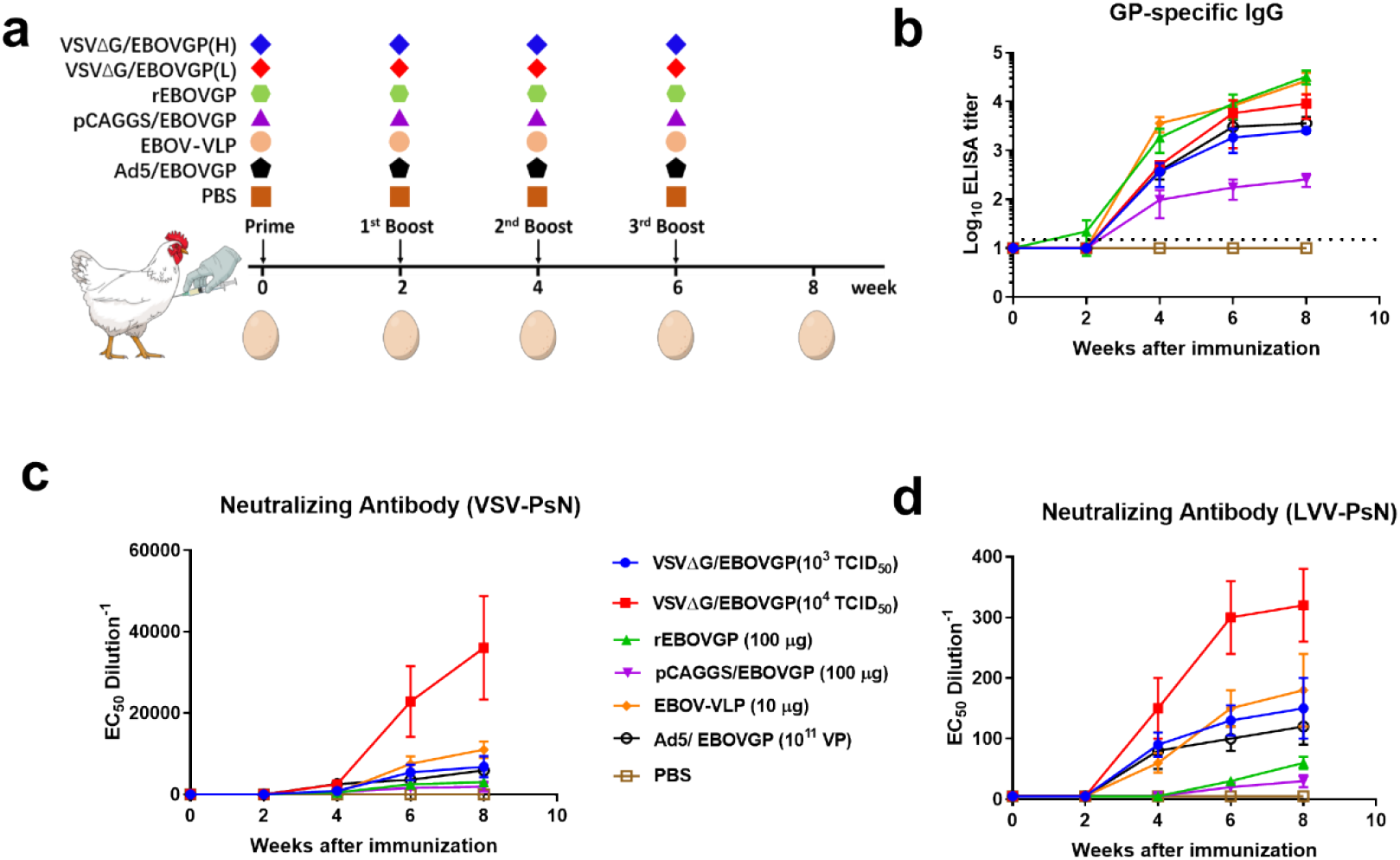
Immunogenicity of different EBOV immunogens in laying hens. (a) Seven groups of laying hens (5-month-old, n=5 per group) were inoculated with VSV Δ G/EBOVGP, rEBOVGP, pCAGGS/EBOVGP, EBOV-VLP, Ad5/EBOVGP or PBS as control, respectively, through i.m. route at 0, 2, 4, 6 weeks. The eggs were collected at 0, 2, 4, 6, 8 weeks. The yolk IgY were purified and used to test the GP-specific ELISA antibody titers (b) and NAb titers (c, d) by VSV-PsN and LVV-PsN assay. The dashed line in B indicates the detection limit. Data are shown as means ± SEM.

### Purification and characterization of IgY

After four immunizations with 10^4^ TCID50 VSVΔG/EBOVGP, IgY antibodies were purified from these collected eggs and then verified by SDS-PAGE (Fig 3a) and Western blot (Fig 3b) assays. SDS-PAGE results showed that two bands with molecular weights of 66 and 22 kDa appeared, representing the heavy chain and light chain of IgY, respectively. The purified IgY antibodies exhibited EBOV GP-specific immunoreactivity to EBOV sGP expressed in *E. coli* expression system when tested with Western blot. Besides, about 50 mg of IgY antibodies with a purity greater than 95% can be obtained from an egg, suggesting that the production cost of this antibody is relatively lower.

**Fig 3.**
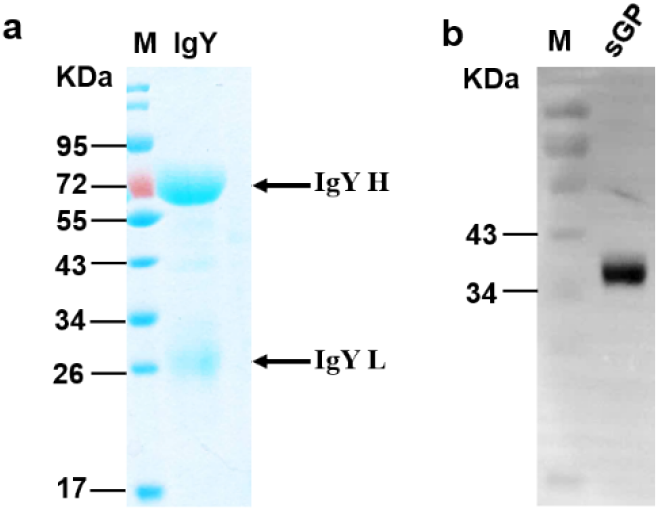
Purity and immunogenicity analysis of anti-EBOV yolk antibody. (a) Purified IgY was analyzed by SDS-PAGE. (b) Recombinant sGP expressed in *E. coli* was analyzed by Western blot with anti-EBOV IgY.

### Passive transfer of IgY protect newborn BALB/c mice from lethal challenge

To determine whether the anti-EBOV IgY antibodies are protective against EBOV, passive protection experiment was performed in newborn BALB/c mice (within the first 3 days of life). Forty newborn BALB/c mice were divided into eight groups, which were challenged subcutaneously (s.c.) with 10^4^ TCID_50_ VSVΔG/EBOVGP. Two hours or 1 day post-infection (dpi), each mouse was adoptively transferred with IgY twice daily for 3 days, and control group mice treated with naive IgY (Fig 4a). To determine the correlation between the transferred IgY dosage and therapeutic efficacy, three different dosages with 10^4^, 10^5^, or 10^6^ NAU/kg (NAb units/kilogram, VSV-PsN) were intraperitoneally (i.p.) transferred to six groups of challenged mice, respectively. The results showed that all the mice receiving 10^6^ NAU/kg IgY were healthy with the gradually increasing bodyweights and without any clinical syndromes within 15 dpi (Fig 4b-c). However, all the control mice died during 6-10 dpi, and the bodyweights began to decrease rapidly from the second day until death. In the other two lower doses administration groups, mice failed to obtain full protection and displayed bodyweight losses after 4 dpi. Only one mouse receiving 10^4^ NAU/kg IgY survived to day 15, and the mice of two timepoints treatment groups increase 5% or 2% of bodyweight with a mean time to death (MTD) of 11.25 or 11 dpi (Fig 4d-e). The situation has improved when the injection amount is 10^5^ NAU/kg. 4/5 mice receiving IgY 2 h post-infection survived to day 15 with an average weight increase of 6% and an MTD of 12 dpi. Another group, 3/5 mice receiving IgY 1 dpi survived to day 15 with an average bodyweight increase of 2% and an MTD of 11.5 dpi. These data indicated that anti-EBOV IgY conferred complete passive protection to newborn mice against a high concentration of VSV vector-based EBOV recombinant chimeric virus challenge. Administrating high-concentration IgY immediately after infection or even one day later can prevent mice from dying by infection, suggesting that anti-EBOV IgY can be used for emergency prevention and treatment after exposure to EBOV.

**Fig 4.**
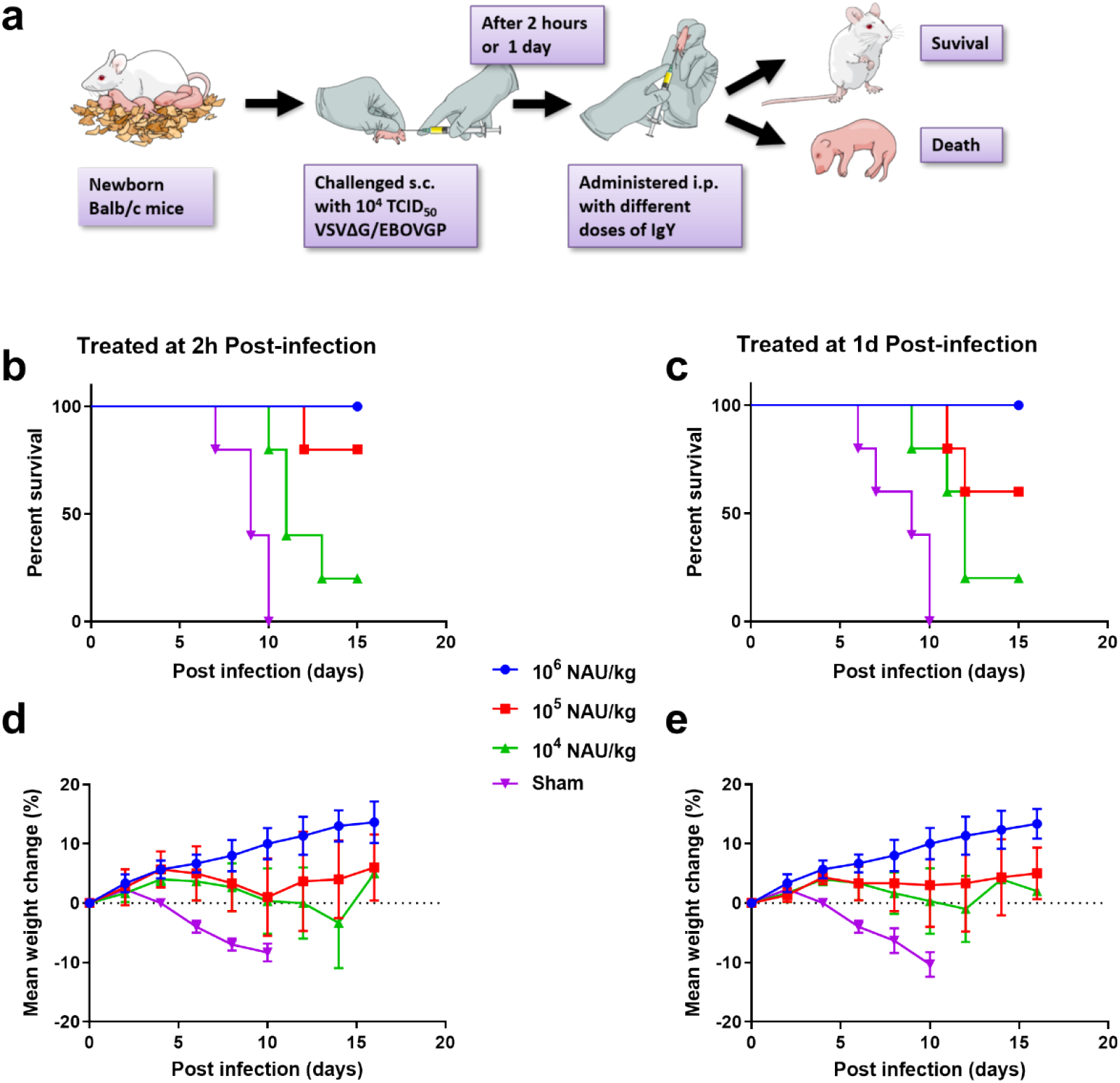
Passive protection of anti-EBOV IgY in newborn Balb/c mice. Experimental scheme (a). Eight groups of newborn Balb/c mice (3-day-old, n=5 per group) were challenged s.c. with 10^4^ TCID50 VSVΔG/EBOVGP, after 2 h or 1 day, each mouse was given different doses of IgY or naïve IgY via i.p. route twice daily for 3 days. The survival rates (b, c) and bodyweight changes (d, e) were monitored every day. Data are shown as means ± SEM.

### Bioavailability of IgY in guinea pigs

To more accurately confirm the protective efficacy of the anti-EBOV IgY, the metabolic level of antibodies in the body needs to be evaluated. The bioavailability of IgY *in vivo* was assessed, two groups of 12-week-old female guinea pigs received s.c. injection with 10^5^ or 10^6^ NAU/kg IgY, respectively. Sera were collected daily and tested by VSV-PsN assay within six days after antibodies injection. The NAbs titer in the guinea pig sera reached a high level. Then the antibody level gradually decreased and fell below the detection limit on the third (low dosage group) and fourth days (high dosage group), respectively (Fig 5). These results suggest that passive transfer of IgY can provide guinea pigs with 2-3 days of protection and that higher doses of antibodies can provide longer protection time.

**Fig 5.**
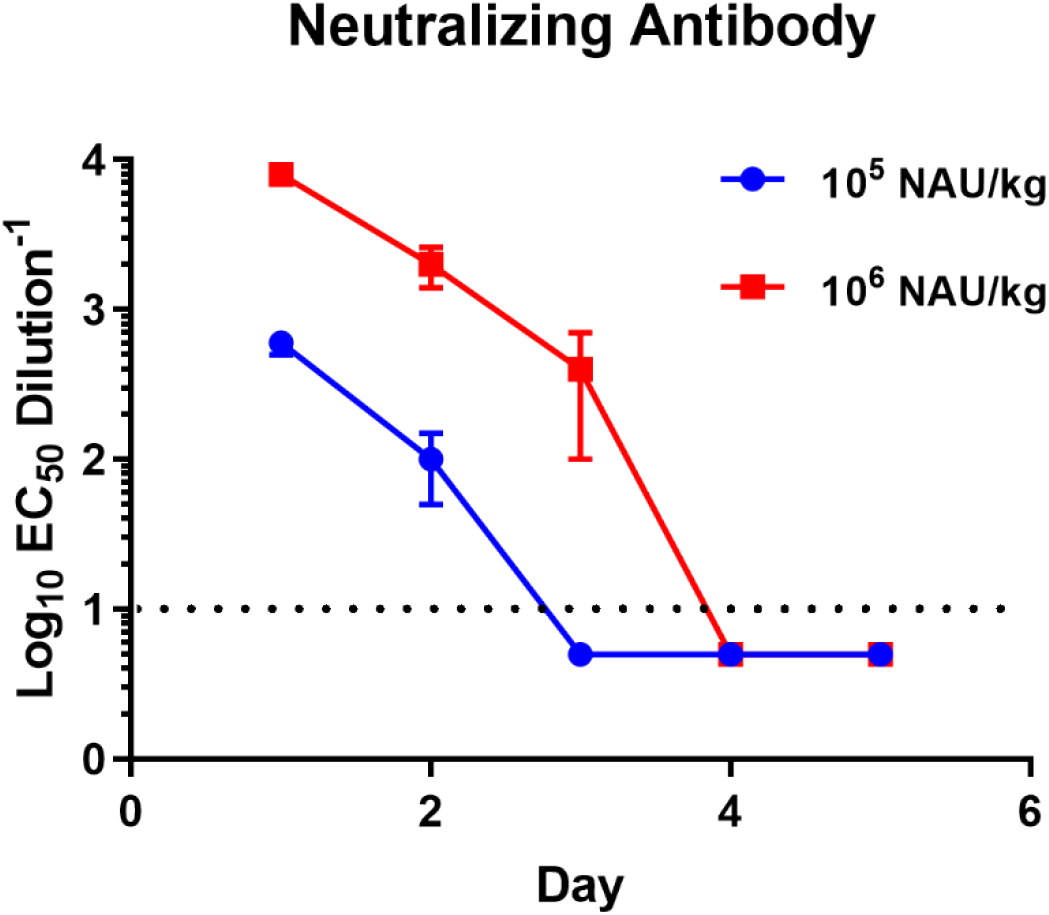
Bioavailability of anti-EBOV IgY in guinea pigs. Two Groups of guinea pigs (12-week-old, n=3 per group) were injected s.c. with 10^6^ NAU/kg or 10^5^ NAU/kg IgY. Sera NAb titers were tested daily by VSV-PsN assay. The dashed line indicates the detection limit. Data are shown as means ± SEM.

### Thermal Stability of IgY

Considering that Ebola hemorrhagic fever mainly occurs in tropical regions, the thermal stability of antibody-based antiviral reagents is essential for practical applications. We stored the purified and filtered sterilized IgY in four different temperature environments, including 4°C, 25°C, 37°C, and 45 °C. Then, the NAb titers were tested every month for one year. The results showed that the IgY NAb titers had no significant change at 4°C and 25°C within one year. However, the antibody titer stored at 37°C gradually decreased from the second month, and only about 20% of the antibody activity remained by the end. In contrast, the activity of the IgY stored at 45°C is lost faster, and the NAbs titer cannot be measured by the third month. These results proved that the anti-EBOV IgY has excellent thermal stability, can be stored at room temperature (RT) for up to one year, and can maintain one month of activity at 37°C without significant changes. Even at a high temperature of 45°C, it still can short-term retention of activity (Fig 6). It is suggested that this anti-EBOV IgY can be used as an emergency prevention/treatment reagent in endemic tropical regions.

**Fig 6:**
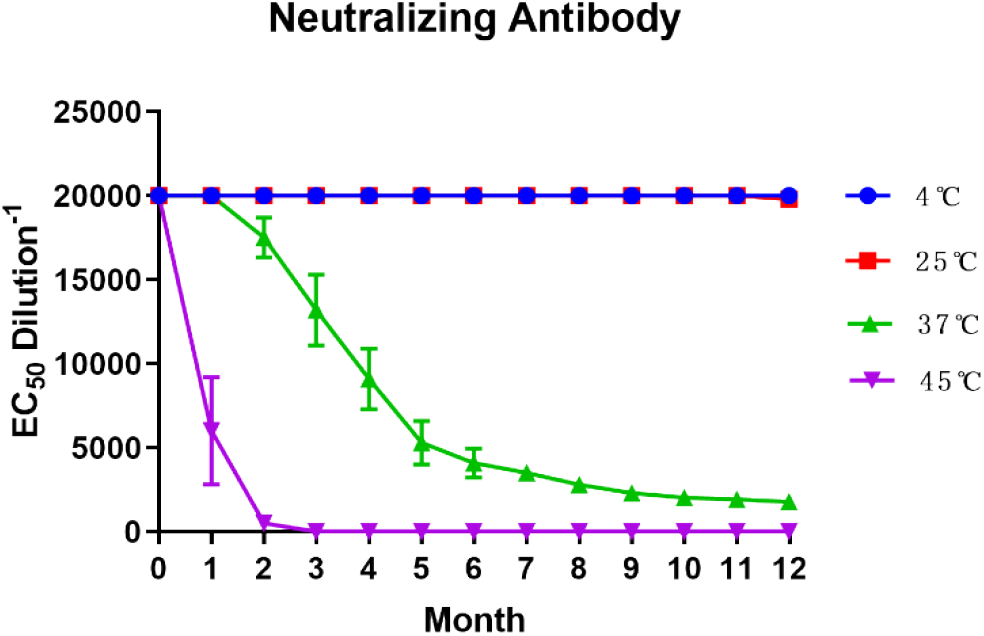
Thermal Stability analysis of purified IgY. Purified IgY were placed under different temperature conditions (4 °C, 25 °C, 37 °C, and 45 °C) for 1 year and neutralizing activity were detected per month.

## Discussion

Ebola hemorrhagic fever is associated with high mortality in humans and spread quickly among populations in which medical facilities are scarce, antibody therapies is the only option for EBOV-infected patients. An ideal therapeutic antibody should be highly efficacious, easy to use, and inexpensive. Previous strategies to develop antibody against EBOV were derived from mammals, although these antibody reagents have proved to have good protective efficacy, their thermal stability is not good [7–11, 23–29]. Considering that EVD mainly occurs in Africa, due to the hot climate and lack of cold chain transportation, the clinical application of these antibodies may be hampered. Moreover, the sooner people receive treatment post exposure to EBOV, the higher the survival rate. Therefore, an Ebola therapeutic antibody with high-efficiency protection and good thermal stability has become a high priority. In recent years, therapeutic antibodies based on avian IgY have begun to receive more and more attention, and a variety of pathogen therapeutic antibodies based on IgY have proved to have very good safety and therapeutic effects [15–19, 21, 22]. Compared with mammalian antibodies, avian antibodies have several distinct advantages. First of all, IgY has higher target specificity, greater binding avidity and longer circulating half-life, which could increase its efficacy against infections [30]; Secondly, IgY does not react with complement system, so the risk of causing inflammation *in vivo* is less. In addition, previous studies have shown that EVD has an ADE effect, which can enhance EBOV infection by human antibodies. Blocking of Fc-receptors reaction through mutating the Fc fragment can abolish ADE, and can effectively enhance the protection efficacy of EBOV antibody [31]. Because IgY does not bind to Fc receptors, so it can perfectly avoid the EBOV ADE effect caused by antibodies. Furthermore, one hen can produce up to 22 grams of IgY antibody a year, which means that the antibody production of two hens is equivalent to one horse, and egg production has already been scaled up, making large-scale and low-cost IgY production feasible [30]. The cheaper antibody drugs are also more suitable for applications in Africa. In addition, compared with antibodies derived from mammalian serum, IgY is derived from eggs, and therefore more in line with animal welfare principles. Finally, IgY has excellent thermal stability, it can even be stored at RT for more than 6 months and at 37°C for more than 1 month, while maintaining its activity [30].

Although multiple EBOV vaccine platforms have been established, several vaccine candidates showed excellent protective efficacy, but a systematic comparison of the humoral immunity levels of different candidates on an animal model has not been addressed. Herein, we tested five different vaccine candidates in laying hens for eliciting yolk IgY level. We demonstrated that the VSV vector-based EBOV vaccine can induce laying hens to produce high-neutralizing IgY, while other vaccines failed to induce a strong humoral immune response. Interestingly, the rVSV vaccine has been widely used (> 40 000 recipients) to prevent the current Ebola outbreak in the DRC [32]. The Ebola vaccine candidates reported earlier have verified its immunogenicity in rodents and primates, but due to the differences in the immune systems of poultry and mammals, these studies cannot be used to direct guide immunization in poultry. The immunogenicity of these vaccines was verified on laying hens, confirming that there is a big difference in the immunogenicity of different vaccines on hens. Fortunately, we obtained an efficient immunization protocol, and prepared high titer neutralizing IgY antibody. To further confirm the protective efficacy of the anti-EBOV IgY, it needs to be verified by the challenge/protection animal model. However, EBOV needs to be operated in the BSL-4 laboratory. Due to limited conditions, we cannot operate live EBOV. We tried to establish an alternative solution. We injected high dose rVSV expressing EBOV GP into newborn mice. We were surprised to find that mice injected with rVSV achieved 100% death, while the control group injected with the same amount of PBS all survival, this phenomenon is limited to mice within 3 days of birth. We injected 1-week-old mice with even higher doses of rVSV and could not cause mice to die, suggesting that the infection model is limited to newborn mice. We used this animal model to evaluate our antibodies and proved that treatment with high-concentration IgY at 2 and 24 hours after challenge can protect mice from death, thus proving that we prepared IgY can protect mice from lethal dose of virus attack.

Antibody therapy played a critical role in antiviral over the past 100 years, humans have exploited passive immunization for treating a variety of infectious diseases. In the West Africa Ebola outbreak, recent evidence suggests therapeutic antibody treatment post-exposure can affect the progression of EVD. The production of recombinantly manufactured MAbs can be clinically useful. Therefore, antibody-based treatments should be further investigated for use in humans. Given the economic and medical restrictions in Africa, all the antibody therapeutic preparations currently have the disadvantages of high cost, long preparation period, and strict need for cold chain transportation conditions. These features suggest there will be an ongoing need for better EBOV therapeutic reagents. Poultry antibodies have the advantages of safety, high efficiency, stability, easy scale production and low cost, which make it very good candidate for global therapeutic use in the time of epidemic.

In this study, we utilized the avian antibodies platform to develop an EBOV therapeutic formulation based on poultry IgY, which is effective when administered as a post-exposure prophylactic in the newborn Balb/c mice model. The conversion of polyclonal antibody products derived from eggs into clinical practice is a daunting challenge. However, ongoing Phase III clinical trials have tested the avian polyclonal IgY against Pseudomonas aeruginosa with the support of the European Medicines Agency. The efficacy of antibodies in the treatment of cystic fibrosis suggests that new methods based on avian antibodies can be used to develop therapies for prevention and treatment. The IgY antibody-based treatments have optimistic research prospects. Compared with other mammalian antibodies, our anti-EBOV IgY antibody has the advantage of greatly improved thermal stability, large quantities lower cost and avoiding ADE reaction, so it is more suitable for African applications. Ebola epidemic areas are in Africa with high temperatures, lack of electricity, and many places do not have cold chain transport and storage conditions. Current mammalian antibodies have limited clinical application due to poor thermal stability. Because of the excellent thermal stability, our developed anti-EBOV IgY provides a promising strategy to solve the current application problems of Ebola antibody-based treatments in Africa.

## Materials and Methods

### Cells and animals

293T, Vero and BHK-21 cells were grown in Dulbecco’s modified Eagle’s medium (DMEM) supplemented with 10% fetal bovine serum and 100 U/mL penicillin-streptomycin (Gibco, Grand Island, NY, USA) at 37°C in 5% CO_2_. Sf9 insect cells were cultured in SF900 serum-free media at 28°C CO_2_-free incubator.

Pregnant Balb/c mice and guinea pigs were purchased from a commercial supplier (Charles River). Laying hens were purchased from Beijing Vital River Laboratory Animal Technology Co., Ltd. All animals were kept in sterile, autoclaved cages and provided sterilized food and water.

### Generation of Immunogens

Five EBOV immunogens based on different platforms were prepared as described previously [33–35], with some modification. All immunogens are designed based on EBOV GP of the 2014 Ebola Makona epidemic strain. Briefly, EBOV GP gene sequence was inserted an additional A residue at position 1019 to 1025 results in a frameshift; thus the complete GP can be expressed. Then the gene is optimized for enhanced transgene expression and then produced synthetically. To obtain DNA vaccine, the GP coding sequence was cloned into a mammalian expression plasmid pCAGGS, transgene expression was verified by western blot. Recombinant EBOVGP (rEBOVGP) protein containing a C-terminal His-tag was obtained through using the insect baculovirus expression system. The EBOV GP gene was first cloned into the pFastBac vector, and the constructed plasmid was transformed into *E.coli* DH10Bac cells to generate a recombinant bacmid. Then Sf9 cells were transfected with recombinant bacmid to generate recombinant baculovirus. After three consecutive passages, the cell culture supernatants were harvested and purified by Ni-NTA affinity chromatography (GE Healthcare, USA). Similar to the preparation of rEBOVGP, to obtain EBOV-VLP, pFastBac Dual vector including EBOV GP and VP40 genes was transformed into DH10Bac, and the resulting bacmid was transfected into Sf9 cells to generate recombinant baculovirus co-expressing EBOV GP and VP40. Culture supernatants were clarified and then pelleted by ultracentrifugation at 30 000 × g for 1 h at 4°C. The pellets were resuspended in PBS and further purified through a 10–50% (w/v) discontinuous sucrose gradient at 25 000 × g for 1.5 h at 4°C. The visible band between 30% and 50% density range was collected and resuspended in PBS. The resulting protein products are EBOV-VLP. A recombinant E1/E3-deleted adenovirus type-5 vector-based EBOV vaccine was created by displacing of adenovirus E1 gene with EBOVGP, forming a recombinant Ad5/EBOVGP genome. The Ad5/EBOVGP was rescued by transfecting the genome into 293T cells and further propagated and purified by CsCl density gradient centrifugation. The number of virus particles (vp) was determined using optical density (260 nm) measurement. The resulting Ad5/EBOVGP cannot replicate inside human tissues. To engineer the recombinant vesicular stomatitis virus (VSV) expressing EBOVGP, the VSVGP gene in rVSV replicon vector pVSV-XN2 was replaced by EBOVGP to generate pVSV-XN2/EBOVGP. Then the recombinant VSVΔ G/EBOVGP was recovered using reverse genetics by co-transfecting pVSV-XN2/ EBOVGP and pBluescript SK+ (pBS) plasmids expressing the VSV nucleocapsid (N), phosphoprotein (P) and large polymerase subunit (L) into BHK-21 cells. rVSV virions were plaque purified, and virus titers were determined by standard plaque assay using BHK-21 cells. All produced EBOV immunogens were verified by western blot.

### Immunizations

Seven groups of 5-month-old laying hens (n =5 per group) were inoculated i.m with immunogens prepared as described above. The detailed scheme is 10^3^ or 10^4^ TCID_50_ VSVΔ G/EBOVGP, 100 μg rEBOVGP, 100 μg pCAGGS/EBOVGP, 10 μg EBOV-VLP, 10^11^ VP Ad5/EBOVGP, or an equivalent volume of PBS as a sham control at weeks 0, 2, 4, 6 (4 times). Eggs were collected at weeks, 0, 2, 4, 6, 8 for ELISA and NAbs test.

### Purification of yolk IgY antibody

IgY was isolated from the egg yolk using the water dilution method, a rapid and simple method was used to separate IgY from the yolk. The separation method is improved based on previous research. Yolks were isolated, removed the yolk membrane, and diluted the contents 1:8 with cold deionized water. Dilution was stirred uniformly, acidified to pH 5.0, keeping stable 10 h then centrifuged at 10 000 × g for 15 min, and the supernatant was gathered. Further purifying the IgY antibody, slowly add 1% (by volume) of the solution of n-octanoic acid with stirring to the supernatant and centrifuge and filter in the same way. Finally, The IgY was further purified by gel filtration (Superdex 200, GE Healthcare), eluted in PBS (pH 7.2) buffer, and concentrated by ultrafiltration to approximately 20 mg/ml using 100-kDa cut-off membranes (Millipore).

### SDS–PAGE and Western blot analysis

Sodium dodecyl sulfate–polyacrylamide gel electrophoresis (SDS–PAGE) was performed to determine the purity of IgY. The samples were mixed with reducing sample buffer, heated at 98 °C for 10 min. Ten microliters of the sample were loaded into each well to SDS-PAGE (12% gel, staining for 3h and destaining for 2h). The pre-stained protein standard (Fermentas, Lithuania) was used as a molecular weight marker. The protein bands were visualized with Coomassie Brilliant Blue R250 (Fluka USA). The gel was analyzed using Bio-Rad image analysis software.

Western blot was performed to check the specificity of the prepared immunogens. VSVΔ G/EBOVGP, rEBOVGP, EBOV-VLP, Ad5/EBOVGP and cell lysate transfected with pCAGGS/EBOVGP plasmid were separated using SDS-PAGE on 15% polyacrylamide gels. For Western blot analysis, the proteins were electrically transferred onto a polyvinylidene difluoride (PVDF) membrane using a semi-dry blotting apparatus (15V, 40 min, RT), then blocked with Tris-buffered saline containing 0.05% Tween 20 (TBS-T) and 5% non-fat dry milk for 1 h at RT and was incubated overnight at 4 °C with a 1:5000 dilution of mouse anti-EBOVGP1 specific MAb. After washed five times with TBS-T, the membrane was incubated with HRP-conjugated goat anti-rabbit IgG diluted 1:2000 (Promega, Madison, Wisconsin, USA) for 2 h at RT. The membrane was washed 4 times. Then specific binding bands were detected by incubation in substrate buffer containing 4 mg 3,3’-diaminobenzidine tetrahydrochloride (Aladdin, China) in 5 mL Tris–HCl and 15 μl hydrogen peroxide for 3–5 min. This reaction was stopped by rinsing with distilled water.

### ELISA

EBOVGP-specific ELISA was used to determine endpoint binding antibody titers of immune yolk IgY. Endpoint titers were defined as the reciprocal serum dilution that yielded an OD450 > 2-fold over background values. Briefly, 96-well plates were coated with 10 μg/mL rEBOVGP in carbonate-bicarbonate buffer, pH 9.6, at 4°C overnight. The plates were then blocked with 5% skim milk in PBS (pH 7.4) at 37°C for 1 h. IgY was added to the top row (1:40), and 2-fold serial dilutions were tested in the remaining rows. The plates were incubated at 37°C for 1 h, followed by five washes with PBST. Subsequently, the plates were incubated with 100 μl of HRP-conjugated rabbit anti-chicken IgY working solution at 37°C for 30 min and washed with PBST five times. The assay was developed using 3,3’,5’,5-Tetramethylbenzidine HRP substrate (TMB) with 100 μl each well stopped by the addition of 50 μl of 2 M H_2_SO_4_ for 10 min. Plates were measured at 450nm by a microplate reader using Softmax Pro 6.0 software (Molecular Devices, CA, USA). PBS was used as a blank control. At the same time, a negative control (IgY derived from PBS-immunized hens) was ascertained in each plate. All ELISA measurements were repeated at least three times with each sample in triplicate.

### Pseudotyped virus neutralization assay

Two pseudoneutralisation (PsN) assays were performed, based on non-replicating VSV pseudotype and lentiviral pseudotype, respectively. EBOV GP pseudotyped lentiviral and VSV virions with a luciferase reporter were produced as previously described [36, 37]. The resulting lentiviral particle can achieve a single-round infection. In brief, Vero cells were plated in 96-well plates and cultured overnight. IgY serial dilutions (1:10, 1:30, 1:90, etc., in DMEM) and pseudoparticles were mixed in a ratio of 1: 9 and incubated at 37 °C for 1 h, before addition to pre-plated target cells in 96-well culture plates (density of 10^4^ cells/well) with 3 replicates. The luciferase activities of infected cells were examined 36 h post-infection. Sample dilutions which showed a 50% reduction in the number of fluorescing cells compared to controls were considered to neutralize antibody titers.

### Adoptive transfer experiment

The yolk was collected and purified two weeks after the final vaccination. Eight groups of newborn BALB/c mice within three days (n=5 per group) were challenged by the s.c. route with 10^4^ TCID_50_ VSVΔG/EBOVGP, after 2 or 24 h, challenged mice were treated i.p. with 10^4^, 10^5^, or 10^6^ NAU/kg anti-EBOV IgY twice daily for 3 days, respectively. Clinical symptoms, bodyweight change rate and survival rate were monitored within 15 days.

### Metabolic studies in guinea pigs

Two Groups of guinea pigs (n=5 per group) were administrated s.c. with 10^6^ or 10^5^ NAU/kg purified anti-EBOV IgY. Sera were collected daily within 6 days for NAbs determination.

## Ethics statement

All animal experiments were approved by the Committee on the Ethics of Animal Experiments of the Institute of Microbiology, Chinese Academy of Sciences (IMCAS), and conducted in compliance with the recommendations in the Guide for the Care and Use of Laboratory Animals of the IMCAS Ethics Committee.

## Acknowledgments

This study was jointly funded the National Key Technologies Research and Development Program of China (No. 2018YFC1200600, 2018YFC1200500) and the National Key Program for Infectious Disease of China (No. 2016ZX10004222-006, 2018ZX10101004, 2018ZX10734-404).

## Author Contributions

Conceived and designed the experiments: YLM LWJ. Performed the experiments: ZY WYQ LYL LY. Analyzed the data: WX TDY JXJ. Contributed reagents/materials: GR. Wrote the paper: YLM ZY.

